# Immune Profiling Panel: a proof of concept study of a new multiplex molecular tool to assess the immune status of critically-ill patients

**DOI:** 10.1101/636522

**Authors:** Dina M. Tawfik, Laurence Vachot, Adeline Bocquet, Fabienne Venet, Thomas Rimmelé, Guillaume Monneret, Sophie Blein, Jesse L. Montogomery, Andrew C. Hemmert, Alexandre Pachot, Virginie Moucadel, Javier Yugueros Marcos, Karen Brengel-Pesce, François Mallet, Julien Textoris

**Affiliations:** EA7426 “Pathophysiology of Injury-Induced Immunosuppression”, PI3, Université Claude Bernard Lyon-1 – Hospices Civils de Lyon – bioMérieux, Lyon, France; Medical Diagnostic Discovery Department (MD3), bioMérieux S.A., France; Immunology Laboratory, Hospices Civils de Lyon, Edouard Herriot Hospital, Lyon, France; Anaesthesia and Critical Care Medicine Department, Hospices Civils de Lyon, Edouard Herriot Hospital, Lyon, France; BioFire^®^ Diagnostics LLC Salt Lake City, UT, USA

**Keywords:** Critically ill patients, Sepsis, Multiplex PCR, Biomarkers, *In Vitro* Diagnostic, FilmArray, Immune response, Syndromic panel

## Abstract

**Background:** Critical illness such as sepsis is a life-threatening syndrome defined as a dysregulated host response to infection and is characterized by patients exhibiting various impaired immune profiles. In the field of diagnosis, a gap still remains in identifying the immune profile of critically-ill patients in the ICU. The availability of an immune profiling tool holds a great potential in providing patients at high risk with more accurate and precise management. In this study, a multiplex immune profiling panel prototype was assessed for its ability to semi-quantify immune markers directly from blood, using the FilmArray® System.

**Results:** The Immune Profiling Panel (IPP) prototype consists of 16 biomarkers that target both the innate and adaptive immune responses, pro- and anti-inflammatory mediators as well as genes involved in diverse regulatory pathways. The analytical studies carried out on healthy volunteers showed minimal inter- and intra-variability in testing the samples across the tested lots. The majority of the assays were linear with an R^2^ higher than 0.8. Results from the IPP pouch were comparable to qPCR and were within the limits of agreement. Finally, quantification cycle values of the target genes were normalized against reference genes to account for the different composition of cells among specimens. The use of the selected panel of markers in IPP demonstrated various gene modulations that could distinctly differentiate three profiles: healthy, borderline mHLA-DR septic shock patients and low mHLA-DR septic shock patients.

**Conclusion:** The Immune Profiling Panel allowed host transcriptomic analysis of immune response biomarkers directly from whole blood in less than an hour. The use of IPP showed great potential for the development of a fully automated, rapid and easy-to-use immune profiling tool, enabling the stratification of critically-ill patients at high risk in the ICU.

## Background

Critically ill patients in the intensive care unit (ICU) exhibit a high risk of morbi-mortality and require special care and timely interventions. One of the major life-threatening situations in the ICU is sepsis, which is defined as an organ dysfunction caused by a dysregulated host response to infection (1). This dysregulated response includes an unbalanced pro- and anti-inflammatory immune response that translates into various immune profiles. These profiles manifest as a state of hyper-inflammation or features of profound immune suppression (2).

This current understanding of the underlying pathophysiology of sepsis has encouraged clinicians to use targeted therapy to restore the immune homeostasis and prevent unfavorable outcomes (3, 4). Nonetheless, personalized care is impeded by the absence of a comprehensive and fast diagnostic tool that would allow clinicians to precisely monitor the patient’s immune profile. For more than 20 years, researchers have described several biomarkers in different platforms to characterize the immune dysfunctions of sepsis (5-8). Transcriptomic gene signatures were identified to stratify septic patients according to the severity and worsening of outcomes that could be used to guide therapy (9-11). Maslove *et al*. sought to validate three proposed scores that can distinguish septic from non-septic patients: Sepsis Metascore (SMS), Septicyte score and FAIM3:PLAC8 ratio in an independent dataset analysis (12). It was shown that the SMS score performed better than the others but further validations are still required. The team endorsed the use of gene expression profiling to stratify patients with sepsis in the presence of a rapid multiplex diagnostic tool (12). However, all the current immune profiling attempts are still in their infancy due to the complexity of the available platforms. Such sophisticated platforms require trained operating personnel, expert bioinformaticians to analyze the data, and experienced clinicians to interpret the results which is highly time-consuming (13). All the previous obstacles hinder the implementation of such technologies as a routine practice in the ICU. However, the recent advances in multiplex-PCR technology could enable the deployment of multiple biomarkers for diagnosing and stratifying patients at the bedside (14, 15).

Multiplex molecular platforms such as the FilmArray® System (BioFire Diagnostics, LLC) have been developed and several commercial kits are available on the market, enabling the accurate detection of pathogens in less than an hour (16). FilmArray is an FDA and CE-IVD certified system, fully automated and user-friendly multiplex-nested qPCR (quantitative Polymerase Chain Reaction) technology that can measure up to 45 assays with a simplified report as a readout (17). We present here a proof of concept study for an Immune Profiling Panel (IPP), a transcriptomic molecular tool assessing the immune status directly from blood. In this work, we report the technical studies of the first IPP prototype used for the semi-quantification of mRNA from blood in the FilmArray System. Finally, the panel was tested on critically ill septic shock patients’ specimens stratified according to the expression of HLA-DR (Human Leukocyte Antigen-DR) on monocytes. A decrease in the expression of HLA-DR on monocytes was often linked to poor outcomes and can be used as a marker to stratify immunocompromised septic patients (18).

## Methods

### 1. Immune profiling panel (IPP)

Several lots of IPP pouch prototypes were manufactured in BioFire® Diagnostics (Salt Lake City, UT, USA) and transferred to our facility for technical assessment. The IPP pouches were run as the commercial syndromic pouches according to manufacturer’s instructions. Briefly, the supplied IPP pouches contain all the biochemical reagents and primers lyophilized ready to use upon hydration, which is done by injecting 1 mL of hydration solution provided with the kit. A 100 μL of whole blood samples were mixed with approximately 800 µL of the lysis buffer provided with the panel and directly injected into the pouch, where a volume of 300 µL of the mix is automatically drawn into the first well (19). Then the pouches were inserted into the FilmArray® 2.0 instrument (BioFire, Inc., Salt Lake City, UT, USA) and nucleic acids were automatically extracted from the sample, then the RNA is reversed transcribed and amplified (17). In some experiments, the extracted RNA samples were tested instead of PAXgene stabilized whole blood to aid in the assessment and study of the panel as some experiments require precise input of RNA. A controlled uniform RNA input helped us to correctly evaluate the semi-quantitative ability of the platform and assess the success of the signal normalization. Since IPP pouches are still a prototype, results are delivered in less than 1 hour in the form of real-time quantification cycle (Cq) values and post-amplification melt peaks. This is different from the commercial kits that provide an easy to read report generated by an internal interpretation algorithm, not yet available in the current IPP prototype.

### 2. Healthy Volunteers and Patients samples

#### Healthy volunteer samples

Whole blood from healthy volunteers collected in PAXgene tubes (Pre-Analytix, Hilden, Germany) was obtained from the EFS (Etablissement Français du Sang, French blood bank, Grenoble). PAXgene tubes were inverted several times and incubated for 2 hours at room temperature according to the manufacturer’s recommendation.

Total RNA was manually extracted from 30 healthy volunteers’ PAXgene stabilized whole blood tubes using PAXgene blood RNA kit (Pre-Analytix, Hilden, Germany) according to manufacturer instructions. The extracted RNA’s quantity and quality were determined using Nanodrop ND-1000 spectrophotometer (Nanodrop Technologies, Wilmington, DE) and Agilent 2100 bioanalyzer (Agilent Technologies, Massy, France) to compute the RNA integrity number (RIN). For the linearity study, 10 extracted RNA with different inputs: 0.5, 1, 2, 10 and 100 ng were directly injected in the amplification chamber of the IPP pouch and samples were run on the FilmArray to study the linearity of the nested PCR assays in the IPP pouch. FilmArray’s IPP performance was compared to qPCR using whole blood and RNA of 30 EFS volunteers. Finally, the extracted RNA of 10 healthy volunteers was tested against septic shock patients’ samples at a quantity input of 10ng.

#### Clinical samples from septic shock patients and cohort details

RNA samples from patients were obtained from a previous prospective study Immunosepsis 1 (IS-1) including adult septic shock patients enrolled from December 2001 to April 2005 from two French university hospital ICUs (20). Twenty septic shock patients’ RNA samples collected on day 3 were selected from the IS-1 cohort according to the expression of mHLA-DR measured by flow cytometry (Table 1). Ten septic shock patients were selected as a low mHLA-DR group when HLA-DR expression on monocytes was less than 30 %. The other 10 septic shock patients had an mHLA-DR expression of more than 30% and were grouped as the borderline mHLA-DR expression group.

**Table 1.** Clinical characteristics of septic shock patients from the Immunosepsis-1 cohort.

| Parameter | Bordeline mHLA-DR Patients (n=10) | Low mHLA-DR Patients (n=10) |
| --- | --- | --- |
| **Characteristics of Patients** |  |  |
| Age (Years) | 68 [47-78] | 66 [36-88] |
| Gender, (Male) | 5 (50) | 5 (50) |
| Comorbidities* ($\geq 1$) | 2 (20) | 5 (50) |
| SOFA (Day 1) | 8.5 [6-14] | 11 [7-15] |
| SOFA (Day 3) | 10 [5-13] | 12 [7-16] |
| SAPS II (Day 1) | 47 [35-67] | 60 [38-89] |
| HLA-DR (% expression on monocytes) Day 3-4 | 56 [35.8-100] | 15.3 [5.2-19.9] |
| **Type of Admission** |  |  |
|  | **N (%)** | **N (%)** |
| Medical | 7 (70) | 5 (50) |
| Elective Surgery | - | 1 (10) |
| Surgical Emergency | 3 (30) | 4 (40) |
| **Primary Site of infection** |  |  |
|  | **N (%)** | **N (%)** |
| Abdominal | 4 (40) | 4 (40) |
| Pulmonary | 4 (40) | 5 (50) |
| Other | 2 (20) | 1 (10) |
| **Outcomes** |  |  |
| ICU length of stay (Days) | 21 [6-82] | 22 [8-47] |
| Survivors at day 28 | 7 (70) | 4 (40) |

Categorical variables are expressed as numbers (percentages), while continuous variables are expressed as median [min-max range].
SOFA: Sequential Organ Failure Assessment score
SAPS II: Simplified Acute Physiology Score II
SAPS II and SOFA scores were measured after 24 hours of ICU stay (Day 1). SOFA score was measured again at Day 3.
\*Comorbidities include: Cardiac, hepatic, respiratory, or/and renal comorbidities

### 3. Reverse transcription and real-time PCR amplification

The qPCR was performed in a microplate, where RNA was reverse transcribed to complementary cDNA using SuperScript® VILO™ cDNA Synthesis Kit (Life Technologies, Chicago, Illinois, USA) and was ready to be amplified. Bench qPCR reactions were performed for S100A9 on a LightCycler 480 instrument (Roche, Switzerland), using its corresponding probes master kit (Roche) following the manufacturer’s instructions. Briefly, PCR reaction was carried out in triplicates in a final volume of 20 μL containing 0.5 μM of primers and 0.1 μM of probe, with an initial denaturation step of 10 mins at 95°C, followed by 45 cycles of a touchdown PCR protocol (10-sec at 95°C, 29-sec annealing at 68-58°C, and 1-sec extension at 72°C). The LightCycler software was used to automatically determine the Cq value for each individual sample and assay. Prototype Argene® kit (bioMerieux, France) was used for the amplification of CD74 and CX3CR1. The kits and RT-PCR amplifications were performed in ABI7500 thermocycler (Applied BioSystems®, USA). Briefly, triplicates of the samples were diluted 1:10 and mixed with 15µl of primer and probe mix, and 0.15µl of RT diluted 1:10 in water to a final volume of 25 µl. The PCR protocol included a 5 mins RT step at 50°C for one cycle, Taq polymerase activation step for 15 mins at 95°C for another cycle. This was followed by PCR protocol of 45 cycles (10-sec denaturation at 95°C, 40-sec annealing at 60°C, and a 25-sec elongation step at 72°C). All samples should give a positive signal at 530 nm (FAM) otherwise the sample is considered negative. Raw Cq values of both methods were evaluated for equivalence using Bland-Altman analysis for each assay individually.

### 4. Statistical analysis & data management

The linearity of markers was evaluated by the visual plotting of the linear regression models of Cq values against the log10 transformation of the RNA quantities and reporting the R squared values (R^2^). Normalized expression values of the genes are expressed as median and interquartile ranges (IQR) box and whisker plots. Paired Wilcoxon signed rank test was used to assess significance before and after normalization of the Cq values in two RNA quantities. The differential expression of the IPP markers between the tested groups was compared using Mann-Whitney U test. The level of significance was set at 5% two-sided tests. Statistical analyses were performed and computed using R software v3.5.1.

## Results

### 1. Immune profiling panel (IPP)

Selection of the IPP markers was based on four pillars: 1) previous laboratory expertise in evaluating the performance and robustness of the markers in clinical trials (21-23) 2) good documentation of prognostic markers in literature (7, 24) 3) performance of the selected assays in duplex and multiplex in a classic qPCR setting 4) address a balanced representation of the pathways involved in diverse cells of both arms of the immunity. The pathways addressed by IPP include both innate and adaptive immune markers that were characterized in previous sepsis studies done in our laboratory. In addition, we aimed to target pro- and anti-inflammatory and immune suppression markers to provide as much information as possible on the immune status and different profiles of patients (Fig.1). Finally, the first prototype of IPP pouches was manufactured which encompassed 16 target assays and 8 reference genes, for the signal normalization. The performance of the markers was then evaluated in several studies.

**Fig. 1.**
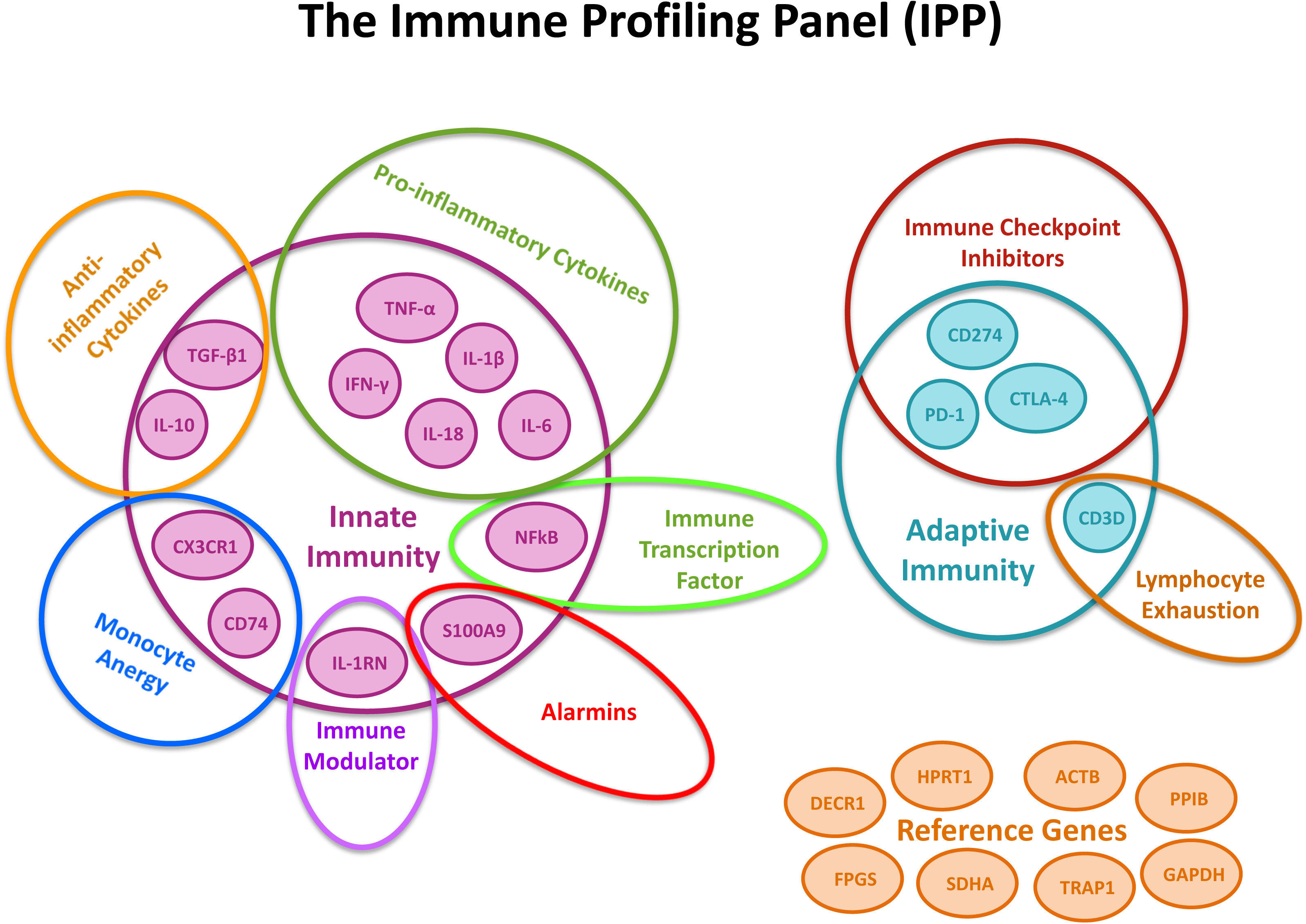
The immune Profiling Panel (IPP). The figure illustrates the selected markers of the panel that includes 16 target genes and describes the different pathways targeted. The panel also features 8 reference genes for signal normalization. The panel of markers was selected to target different arms of the immune responses (innate and adaptive), several immune functions (pro- and anti-inflammatory cytokines) and immune pathways.

### 2. Repeatability study

We inspected the repeatability among 4 manufactured prototype lots by testing whole blood collected from a single healthy donor tested in triplicates. The variance was computed for all the samples and assays in each lot. A threshold of variance acceptance was set to +1 SD (Standard Deviation) of the overall variance. This threshold was selected as it was more stringent compared to the usually recommended +2 SD and could identify the markers with high variance. Fig. 2 illustrates that the overall variance for all markers was low across lots. Lot B had a higher variance which was mainly due to one gene (SDHA, a reference gene) that seems to be also variable in lot D. CD74 was identified as an outlier in one occurrence in only lot B while, PD-1 was an outlier in lots C and D. The observed high variances seem to be assay-related rather than lot-related. Indeed in the case of SDHA, the variance is probably linked to a problem in primer design that can be also be observed in the following studies, while PD-1 variability might be due to the fact that it is barely expressed in healthy volunteers. The rest of the assays had minimal variability and remain below the limit of +1 SD, demonstrating the repeatability and robustness of the system.

**Fig. 2.**
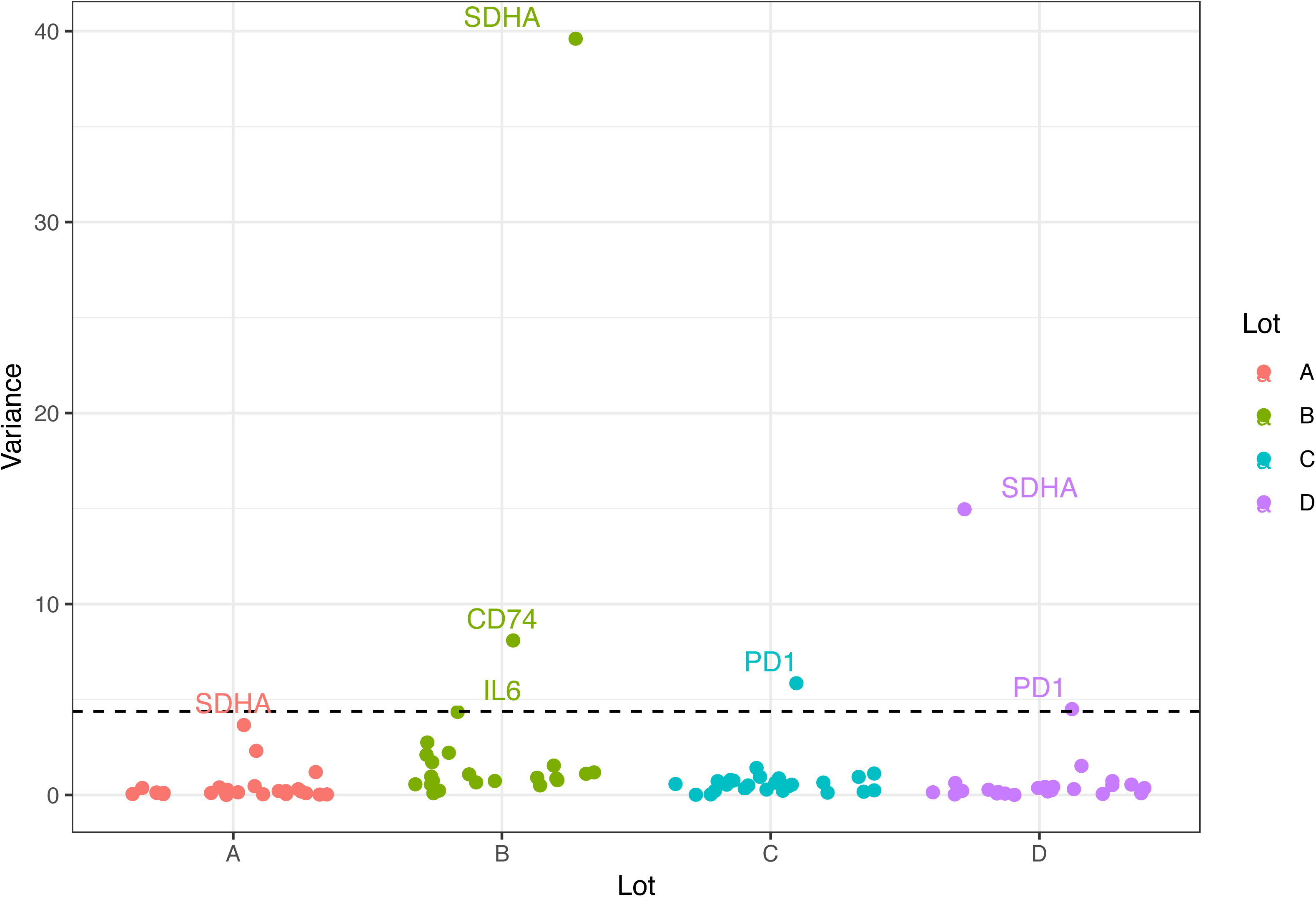
PAXgene stabilized whole blood from a single healthy donor was tested in triplicates to evaluate the variability of the assay in the IPP tool. The variance in Cq values of 4 different lots given as A, B C and D are presented on the y-axis calculated from the triplicates of the markers expression across each lot with a cut-off + 1 SD. The name of the target genes above or on the line of variance cut-off are indicated in the plot.

### 3. Linear study of IPP assays in FilmArray

The FilmArray platform was initially developed for microbiology applications and detection of various pathogens from different sample types. Montgomery *et al*. initiated a study to use host response-based assays in FilmArray to discriminate viral from bacterial infection in patients (25). In our prototype pouch, we wanted to semi-quantify the host immune biomarkers in critically ill patients. To this end, we studied the linearity of the selected assays to ensure the possibility of semi-quantification using the FilmArray system. We used 5 known RNA quantities (0.5 – 100ng) to show that the IPP markers expression fall into the tested linear range of measurement. Fig.3 illustrates the linearity of three reference genes (DECR1, HPRT1, and PPIB) and three target genes (S100A9, CD74 and CX3CR1) representative of the panel. Reference and target gene assays were linear within the tested range of the total RNA quantities with R^2^ values ranging from 0.51 to 0.94 (median 0.89). The majority of IPP assays exhibited high R^2^ values above 0.8 (Table 2). R^2^ values below 0.8 were further inspected; such as IL6 and PD-1 which might have a low R^2^ due to their weak expression in healthy volunteers as they are prominently expressed in ill patients. Genes such as GAPDH, SDHA and ACTB showed poor performance and were discarded from the rest of the analytical studies as they were not linearly expressed with different RNA quantities. Overall, this confirms that FilmArray’s IPP assays are linear enabling semi-quantification of genes’ expression using RNA extracted from whole blood samples.

**Fig. 3.**
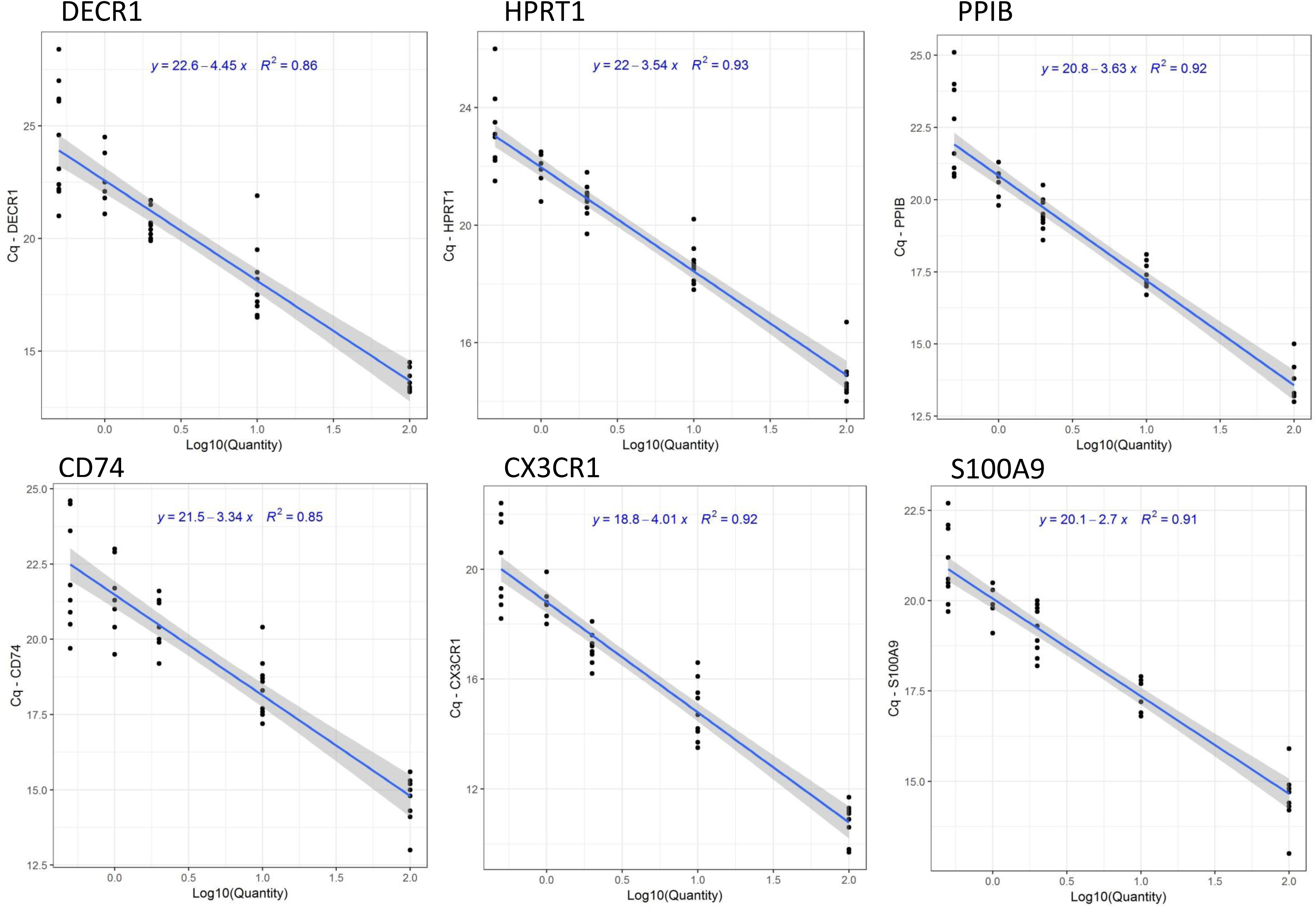
Linearity study of reference and target assays. Extracted RNA from 10 healthy volunteers was tested in IPP using 5 different RNA quantities (0.5-100 ng). The linear model of the Raw Cq values is plotted against log 10 of the RNA quantities. The slope-intercept equation of each model appears on the plot along with the R^2^ values.

**Table 2.** The global R^2^ values of the Immune Profiling Panel (IPP) markers.

| A. Target Marker | $R^2$ | B. Reference Marker | $R^2$ |
| --- | --- | --- | --- |
| IL6 | 0.51 | ACTB | 0.63 |
| PD1 | 0.63 | SDHA | 0.74 |
| IL10 | 0.80 | GAPDH | 0.78 |
| IL1RN | 0.82 | DECR1 | 0.86 |
| TGFB1 | 0.85 | TRAP1 | 0.89 |
| CD74 | 0.85 | PPIB | 0.92 |
| IFNg | 0.86 | HPRT1 | 0.93 |
| IL1B | 0.87 | FPGS | 0.94 |
| CD3D | 0.87 |  |  |
| IL18 | 0.88 |  |  |
| NFkB | 0.88 |  |  |
| CTLA4 | 0.89 |  |  |
| S100A9 | 0.91 |  |  |
| CD274 | 0.91 |  |  |
| CX3CR1 | 0.92 |  |  |
| TNFa | 0.93 |  |  |

### 4. Equivalence to qPCR

Three target genes were tested (S100A9, CD74 & CX3CR1) for equivalence between the two methods using Bland-Altman plots (Fig. 4). It was observed that all the three assays are within the limits of agreement demonstrated as ± 1.96 SD calculated from the mean difference horizontal line. S100A9 and CD74 were equivalent in both platforms as most of the points are around the mean difference line on the y-axis which is close to zero. Whereas in qPCR, CX3CR1 presents a higher systematic bias of 5.9 Cq compared to IPP and a slight decreasing proportional bias associated with higher Cq values, but it still remains within the limits of agreement (Fig. 4A). After normalization of both data and re-computing the Bland-Altman plots, it can be observed that normalization helped eliminate the proportional bias with a slight presence of a systemic bias between the two methods for the three markers (Fig. 4B). Both raw and normalized Cq analyses show that the two methods are within the limits of agreement. This analysis demonstrates the concordance between FilmArray’s IPP and bench PCR which is a common reference method used for mRNA quantitation.

**Fig. 4.**
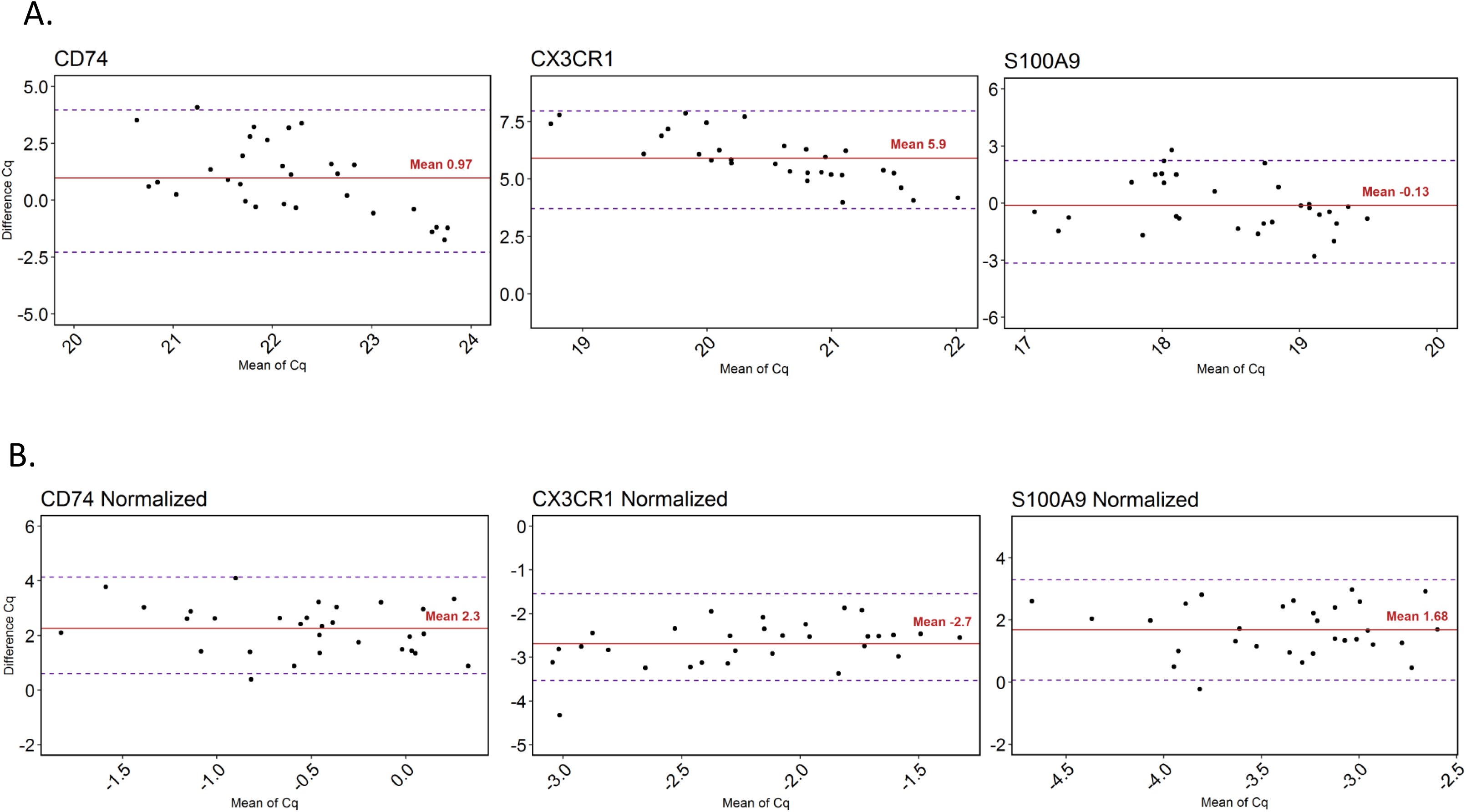
Bland-Altman analysis of 3 target markers expression in the IPP pouch compared to qPCR results. Whole blood from PAXgene tubes of 30 healthy volunteers samples was tested in FilmArray and extracted RNA samples from the same volunteers was tested in qPCR for the equivalence study. The red horizontal line represents the mean difference and estimates the systemic bias between the methods, the dispersion of points is enclosed within ± 1.96 SD limits of agreement is presented as dashed lines. All three genes are within the limits of agreement, A. Raw data B. Normalized.

### 5. Evaluation of FilmArray’s IPP Signal Normalization

Normalization is critical for the signal correction as the assays are tested in a fixed input volume of whole blood with an unknown quantity of RNA. Ten healthy volunteers were further analyzed to assess the effectiveness of the normalization strategy. Two RNA inputs (2 ng and 10 ng) were tested and the expression signal was represented in boxplots before and after normalization. Fig. 5 illustrates 3 target genes (CD74, CX3CR1, and S100A9 selected as representative of the data). Fig. 5A shows a significant difference in expression level (p < 0.01) presented by the raw Cq values that are dependent on the RNA quantity (as previously inspected in the linearity study). Fig. 5B shows the same data after normalization with the internal reference genes. The results in Fig. 5B shows that after normalization the different RNA inputs were corrected and medians were not different.

**Fig. 5.**
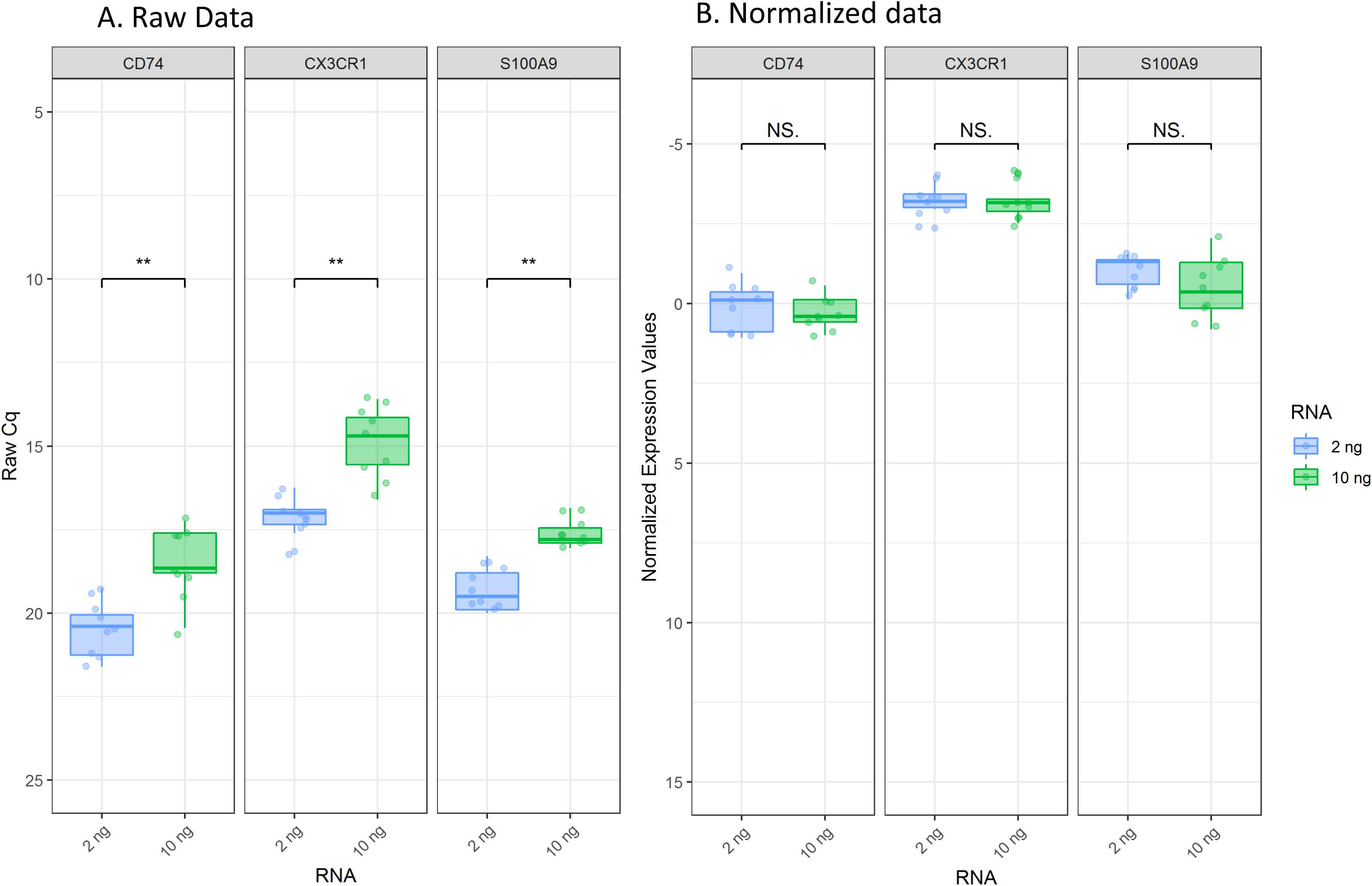
Evaluation of Data normalization: The raw Cq values and normalized expression values are expressed in an inverted y-axis to facilitate interpretation. RNA samples from 10 healthy volunteers extracted from PAXgene tubes were directly injected in the IPP pouch in two different quantities and tested in FilmArray. The results were tested for significance using paired Wilcoxon signed rank test (where * is p < 0.05 and NS as not-significant). A. Shows the raw Cq values of 10 healthy volunteers expressed as boxplots showing the 2 RNA quantities tested; 2 (blue) and 10 (green) ng. B. Shows the boxplots after Cq normalization with no significant difference observed between 2 quantities in the respective marker.

### 6. Clinical samples testing with IPP

To ensure that the expected intra-variable expression of markers between patients with different immune status and healthy volunteers are conserved after normalization, we ran a proof of concept analysis on 10 healthy volunteers against 20 septic shock patients stratified using mHLA-DR. Testing the samples with IPP showed differential expression of the target genes across the 3 tested populations. Fig. 6A shows 6 genes that were down-modulated in patients compared to healthy volunteers. A significant difference between the patients and the healthy group was observed in CD74, CX3CR1, CD3D, CTLA-4 and IFN-⍰. These assays cover diverse immune functions and are characteristics of monocyte anergy, lymphocyte exhaustion, antigen presentation, and pro-inflammatory cytokine production. All the previous dysfunctions and modulations are hallmarks of sepsis and can be clearly observed in both septic shock groups, the borderline and the low mHLA-DR expression patients, with more aggravation in the latter group. Fig. 6B shows 4 assays that were significantly up-modulated in patients that include IL-18, IL-10, IL1RN and S100A9. These markers are related to pro- and anti-inflammatory cytokines and danger associated molecular pattern (DAMPs) alarmins related to both arms of the immune response (innate and adaptive). Interestingly, the use of the stratified samples showed the ability of the IPP tool to clearly distinguish between healthy volunteers and patients with a various degree of immune-alterations, that specifically identified the immunosuppressed profiles among the septic shock patients.

**Fig. 6.**
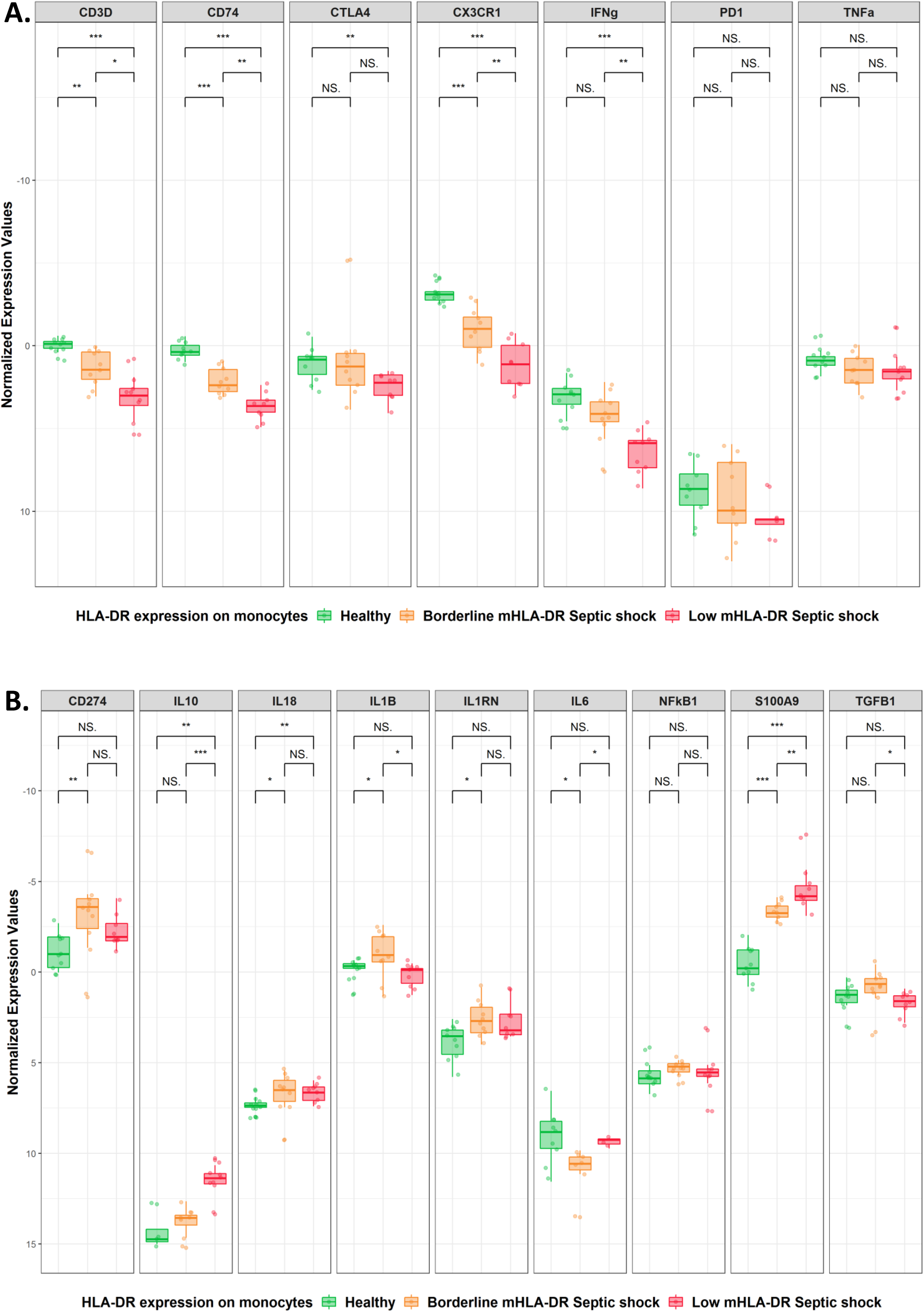
Testing the immune profiling panel on 10 healthy volunteers and 20 septic shock patients: 10 in the borderline mHLA-DR group and 10 in the low mHLA-DR group. The y-axes representing normalized values are inverted to facilitate interpretation. A quantity of 10ng of each RNA sample was injected directly in the IPP pouches. Normalized expression values were compared across groups using Mann-Whitney U test for significance. **A.** Shows the markers that were down-modulated in the patient groups compared to the healthy volunteers. **B.** Illustrates markers that were up-modulated in patients against healthy volunteers (where NS: p > 0.05, *: p < 0.05, **: p < 0.01 and ***: p < 0.001)

## Discussion

A proof of concept study was set up to assess the IPP’s ability to semi-quantify immune-related markers on the technical and clinical levels using samples from both healthy volunteers and patients. The selected panel was able to differentiate between the healthy volunteers and the two groups of septic shock patients. Our results suggest the potential use of IPP as a tool to stratify patients according to their immune status.

The recent advances in diagnostic techniques such as multiplexing PCR, bead-based proteomics and cell phenotyping panels approaches are now shaping the landscape of patient care and advances towards precision medicine (14). As previously reported in the literature, patients in the ICU, especially septic patients, are heterogeneous and their immune response is highly dynamic. However, patient management in the ICU remains a “bundle of care” with only a slight modification according to the clinicians’ experience rather than personalized care (4, 26). To reduce this diagnostic gap in detecting impaired immune responses, many research efforts sought after immune profiling approaches to stratify patients at risk (13, 27). To develop a comprehensive immune profiling tool, a panel of biomarkers should be considered to cover various immune functions as described previously, including the innate, adaptive, pro- and anti-inflammatory immune responses. Having such tool as a point of care will probably help clinicians achieve patient-guided management and therapies.

In this work, we present the first proof of concept on the Immune Profiling Panel (IPP), a new multiplex molecular tool to assess critically-ill patients’ immune status in the FilmArray System. FilmArray’s microbiology syndromic panels have been reported in the literature for their robust and reproducible results but they mainly provide qualitative results for pathogen detection (16, 17, 28). In the IPP tool, a different approach was sought, and we demonstrated the ability of the platform to semi-quantify immune host-response markers. Even though the capacity of the pouch reaches up to 45 assays, we chose to use only 16 target genes and test 8 reference genes (Fig. 1) as a prototype. Developments are currently undergoing to achieve a higher multiplexing capacity of the IPP tool.

The IPP pouches were analytically evaluated for repeatability, for assay’s linearity and was compared to qPCR which is a well-established gold standard for analyzing the transcriptome from whole blood. As explained by the MIQE guidelines for qPCR validation, repeatability is the intra-assay variation reported by SD or variance in Cq values (29). Most of the assays had a low variance, except for only SDHA which was highly variable (Fig. 2) and recorded the highest coefficient variance of 19 % (data not shown), and was thus discarded later. All the assays were further investigated in the linearity study (Fig. 3) in a 3-log linear range of RNA inputs. We confirmed that the majority of the markers in the panel had an R^2^ ranging from 0.8 – 0.95. These values are slightly lower than the recommended R^2^ (≥0.98) for classic quantitative PCR (30). However, the qPCR recommendations only address a one-step amplification PCR whereas the R^2^ presented here covers the whole process from sample input to result. Equivalence studies to classic qPCR showed that the assay results were concordant with assays from IPP with a low bias for CX3CR1 (Fig.4). The observed R^2^ and bias might be explained by several reasons such as the integration of several steps in the platform that include reverse transcription, multiplex amplifications that is followed by a dilution step just before the second round of nested PCR that might slightly influence the quantity of RNA and the signal at the end of the run. Other factors might be the high multiplexing capacity of the platform as some primers might interact and affect one another within the multiplexing environment, in addition to the natural variability in expression profiles of healthy volunteer samples obtained from the blood bank. All these factors make the standard guidelines more adaptable to classic qPCR and are partially applicable to the FilmArray System assessment. Nevertheless, the achieved linearity is acceptable and allows assays’ semi-quantification of mRNA from whole blood. Based on the analyses, the following assays SDHA and ACTB were discarded and were not included in the later assessment steps as they likely have a design or compatibility issues. The rest of the assays that were identified in the variability study or had an R^2^ lower than 0.8 were investigated and will either re-designed or removed from the next version of the tool. Finally, since the intended test specimen or matrix is whole blood, the effectiveness of signal normalization was confirmed by the successful correction of the varying RNA input among individuals that can influence the RNA quantity within the sample (Fig. 5).

A decrease in mHLA-DR expression measured by flow cytometry is widely accepted as a marker of immune suppression in critically-ill patients. mHLA-DR expression lower than 30 % was often associated with mortality and risk of developing secondary infections at day 3-4 after sepsis onset (18, 31). When IPP was tested on healthy and septic shock patients (Fig. 6), differential expression was shown and a basic stratification of patients was possible according to their immune profile. For instance, the panel successfully pointed out that low mHLA-DR patients suffered more profound immune dysfunction than the borderline mHLA-DR patients. This was highlighted by the down-modulation of CD74 and CX3CR1, CD3D and CTLA-4 markers that are affiliated to the innate and adaptive immune responses, respectively, compared to healthy subjects. Other immune dysfunctions that were observed include alterations in both pro- and anti-inflammatory markers (IL-18, IL-10 and IFN-⍰). These markers are reported in the literature as hallmarks of sepsis syndrome and are indicators of an immunosuppressed profile in septic shock patients (2). The importance of having an immune profiling panel lies in the valuable information provided by the tool about several aspects of the immune response, dysfunctions and physiopathologies that cannot be identified by measuring only one aspect or a unique maker such as HLA-DR.

Recent studies were proposed by researchers to overcome the diagnostic gap in immune dysfunction profiling using different platforms. For instance, a microfluidic biochip has been developed based on the quantification of CD64 from circulating neutrophils in the blood and enumerating the lymphocyte count using only 10 µl of blood from patients in 30 minutes. The microfluidic chip technology could potentially stratify sepsis in the patient population (32). This approach is rather appealing but due to the heterogeneity of immune responses in sepsis patients, both the diversity and number of the addressed biomarkers become key in the precise stratification, diagnosis and prognosis. Similarly, interesting work by Morris *et al*. based on a 4-hours flow cytometry protocol assessing neutrophil CD88, percentage of regulatory T cells (T_regs_), and mHLA-DR expression demonstrated the potential to predict secondary infections in septic patients (6). However, the main challenges that hinder the use of flow cytometry at the bedside, remains that it is mainly operated and interpreted by skilled personnel which makes it hard to standardize, and requires the presence of well-equipped laboratories that work round the clock which is not the case in most hospitals. These research efforts re-enforce the need for a tool such as IPP in the ICU, as it includes a panel of markers that cover diverse immune functions and can identify different patient profiles. The fact that FilmArray is a fully automated and closed system with only 2 minutes of hands-on time, limits the risk of variability and facilitates its implementation. The use of whole blood as an input and the availability of results within the hour makes it possible to be installed at the central laboratory or at the bedside, thus making it accessible 24/7. Our proof of concept provides great promise to apply molecular multiplexing technology in immune profiling of critically-ill patients as a point of care in the ICU.

In this pilot study, we have several limitations such as the small sample size of patients, as most of our technical evaluations were on healthy volunteers. In addition, markers performance such as validity and ability to predict clinical outcomes still need to be addressed in a dedicated clinical cohort. The further addition of assays in next pouch versions will require a full analytical validation and evaluating the compatibility of all primers in the multiplexing environment. Our upcoming goal is to increase the multiplexing capacity of the panel in such a way to have a highly informative tool reflecting the immune status of a patient at a given time. The final panel will be more comprehensive, having a simplified readout that can be integrated into a day-to-day clinical practice in the ICU. This will provide personalized information for each patient and will enable clinicians to precisely manage critically-ill and sepsis patients according to their immune profile.

## Conclusion

The Immune Profiling Panel is a new molecular multiplex tool that uses the FilmArray System which provides a transcriptomic immune profile of critically ill patients in the ICU. The analytical assessment proved the ability of the selected panel to measure immune-related gene expression from blood in both healthy and septic shock patients. The easiness of use and rapid time-to-result of IPP proves its great potential for the development of a full capacity and automated tool to be used at the bedside. The IPP tool could be used in the future to monitor and stratify patients at high risk of secondary infections and mortality, based on their immune status, enabling personalized patient care.

## Abbreviations

IPP: Immune Profiling Panel
ICU: Intensive Care Unit
mHLA-DR: monocytic Human Leukocyte Antigen-DR
Cq: quantification cycle
RT: Reverse Transcription
qPCR: quantitative polymerase chain reaction
RNA: RiboNucleic Acid
SD: Standard Deviation

## Acknowledgements

We acknowledge Estelle Peronnet, Elisabeth Cerrato and Boris Meunier for their technical support and sharing their expertise to enrich our study. We extend our gratitude to the Immunospesis-1 study group for providing us with the septic shock clinical samples.

## Declarations

### Ethics approval and consent to participate

Healthy volunteers: Whole blood from healthy volunteers collected in PAXgene tubes was obtained from the EFS (Etablissement Français du Sang, French blood bank, Grenoble). Informed consents from the blood donors were obtained and their personal data were anonymized at time of blood donation and before the blood transfer to our facility according to EFS standard regulations for blood donation.

Septic shock patients: Immunospesis-1 cohort was approved by the local ethics committee (Comité de Protection des Personnes Sud-Est II #IRB 11236). A non-opposition to cohort inclusion was recorded from every patient or the patient’s relative. Since it was a non-interventional trial and complementary blood samples were obtained during patients’ routine blood sampling and tested after the completion of routine follow-up tests, no informed consent was required. The Immunosepsis-1 study is registered at the French Ministry of Research and Teaching (#DC-2008-509), at the Commission Nationale de l’Informatique et des Libertés and on clinicaltrials.gov (NCT02803346).

### Consent for publication

Not applicable.

### Availability of data and materials

Kindly contact the corresponding author for the data. Data can be provided upon motivated requests.

### Authors’ contributions

All authors contributed to the manuscript structuring and critical revision for important intellectual content, read and approved the final manuscript. LV, AB, FM, JYM and JT designed the experiments. DMT and AB performed the experiments and the statistical analyses. DMT drafted the manuscript. DMT, LV and JT made substantial contributions to the conception and design of the study and final data interpretation.

